# GalaxyCloudRunner: enhancing scalable computing for Galaxy

**DOI:** 10.1101/2020.05.28.121772

**Authors:** N Goonasekera, A Mahmoud, J Chilton, E Afgan

## Abstract

**Summary:** The existence of more than 100 public Galaxy servers with service quotas is indicative of the need for an increased availability of compute resources for Galaxy to use. The GalaxyCloudRunner enables a Galaxy server to easily expand its available compute capacity by sending user jobs to cloud resources. User jobs are routed to the acquired resources based on a set of configurable rules and the resources can be dynamically acquired from any of 4 popular cloud providers (AWS, Azure, GCP, or OpenStack) in an automated fashion.

**Availability and implementation:** GalaxyCloudRunner is implemented in Python and leverages Docker containers. The source code is MIT licensed and available at https://github.com/cloudve/galaxycloudrunner. The documentation is available at http://gcr.cloudve.org/.

**Contact:** Enis Afgan

**Supplementary information:** None

## 1. Introduction

Galaxy (Afgan, Baker, et al., 2018) is a popular data analysis and tool integration platform used in a variety of research scenarios, with public Galaxy servers, such as the *usegalaxy*.* federation (Afgan, Jalili, et al., 2018), offering free resources for scientific analyses. However, batches of genomic sequencing data, workshop training events, or specific resource requirements (e.g., large memory) are examples of scenarios where additional compute resources can deliver more responsive and suitable environments for users. For such scenarios, public servers are challenging to use due to the limited processing capacity often leading to prolonged job wait times (Tyryshkina et al., 2019). Meanwhile, setting up and maintaining suitable and sizable resources locally presents challenges such as acquiring the necessary infrastructure, which may often be underutilized.

Cloud computing offers opportunities to acquire suitable resources on demand (Langmead and Nellore, 2018). However, behind a seemingly straightforward Galaxy web interface used for job submission, the Galaxy application performs a large number of steps when executing jobs. These include ensuring the availability of necessary tools, formatting job submission scripts, staging job input data, submitting and monitoring jobs, and retrieving outputs. Ensuring these steps are properly configured imposes significant complexity on the system administrator when trying to leverage cloud resources (Afgan *et al.*, 2015).

To this end, GalaxyCloudRunner automates the necessary steps for Galaxy to provision, connect, and route jobs to remote machines. Based on configurable and extensible rules, a Galaxy administrator can enable cloud bursting for their Galaxy instance, as well as have control over which jobs are sent to remote resources, how many and which machines are made available, and to mix and match which clouds they are running on. Meanwhile, the end-user seamlessly uses Galaxy while benefiting from the increased compute capacity.

## 2. Implementation

The GalaxyCloudRunner is implemented as a Galaxy plugin and comes built into Galaxy versions >=19.01. By leveraging Pulsar, Galaxy’s remote job runner, and CloudLaunch (Afgan *et al.*, 2019), a cloud resource launcher, GalaxyCloudRunner assembles all the necessary components to enable cloud-busting, while providing configurable dynamic job routing rules. Together, GalaxyCloudRunner, Pulsar, and CloudLaunch allow for the dynamic creation of virtual machines (VMs) on any of popular cloud providers, namely Amazon Web Services, Google Cloud Platform, Microsoft Azure, and OpenStack. The provisioned VMs are automatically configured to accept jobs from a Galaxy instance, with user jobs being routed following desired rules.

GalaxyCloudRunner operates in two distinct steps: (1) a Galaxy administrator enables GalaxyCloudRunner as a job destination by editing a single Galaxy configuration file, which defines job rules (more below) and a connection to CloudLaunch, and (2) the administrator launches VMs via CloudLaunch, which can be done manually via a graphical web interface or programmatically. A public instance of CloudLaunch is available at https://launch.usegalaxy.org/. Galaxy will continuously query CloudLaunch for available machines. Each time, CloudLaunch will return the list of available Pulsar servers along with tokens that Galaxy can use to authenticate with each Pulsar server. This allows the administrator to dynamically add or remove machines as desired, which can be scripted for a hands-off solution. Based on the availability of machines and following the defined set of rules, Galaxy will route jobs to them.

The GalaxyCloudRunner implements a number of job rules, which can be chained or extended for custom routing capabilities. The default rule (`galaxycloudrunner`) has Galaxy querying CloudLaunch for any VMs available to receive jobs, which are then dispatched in round-robin fashion. Additional rules are available to limit sending jobs to the `galaxycloudrunner` only when the main job queue has a backlog. Such rules allow the utilization of local resources first, and only bursting to the cloud when necessary. A rule also exists to route jobs based on input size, such as checking if inputs are too large for local machines, or conversely routing numerous small jobs to the cloud that do not require large data transfers. These existing rules can be chained together to enable behavior such as sending all jobs smaller than 1GB to remote resources when the local queue has a backlog greater than 5 for example. Additional rules can be developed by administrators to implement any desired logic for job routing.

When an administrator launches a VM via CloudLaunch, CloudLaunch ensures a suitable runtime environment exists on the remote machine by automatically starting Pulsar and configuring Galaxy CVMFS, a read-only filesystem repository managed by the Galaxy project and containing a large number of pre-installed tools and reference data (Afgan, Baker, *et al.*, 2018). The Pulsar server uses Slurm as a job manager, and will queue jobs if the machine lacks sufficient processing capabilities for the current load. At job submission, Pulsar stages input data and retrieves tool binaries from the CVMFS repository. If a tool is unavailable on CVMFS, Pulsar will attempt to install it via Bioconda recipes (Grüning *et al.*, 2018). Once a job completes, results are automatically transferred back to Galaxy (Figure 1).

**Fig. 1.**
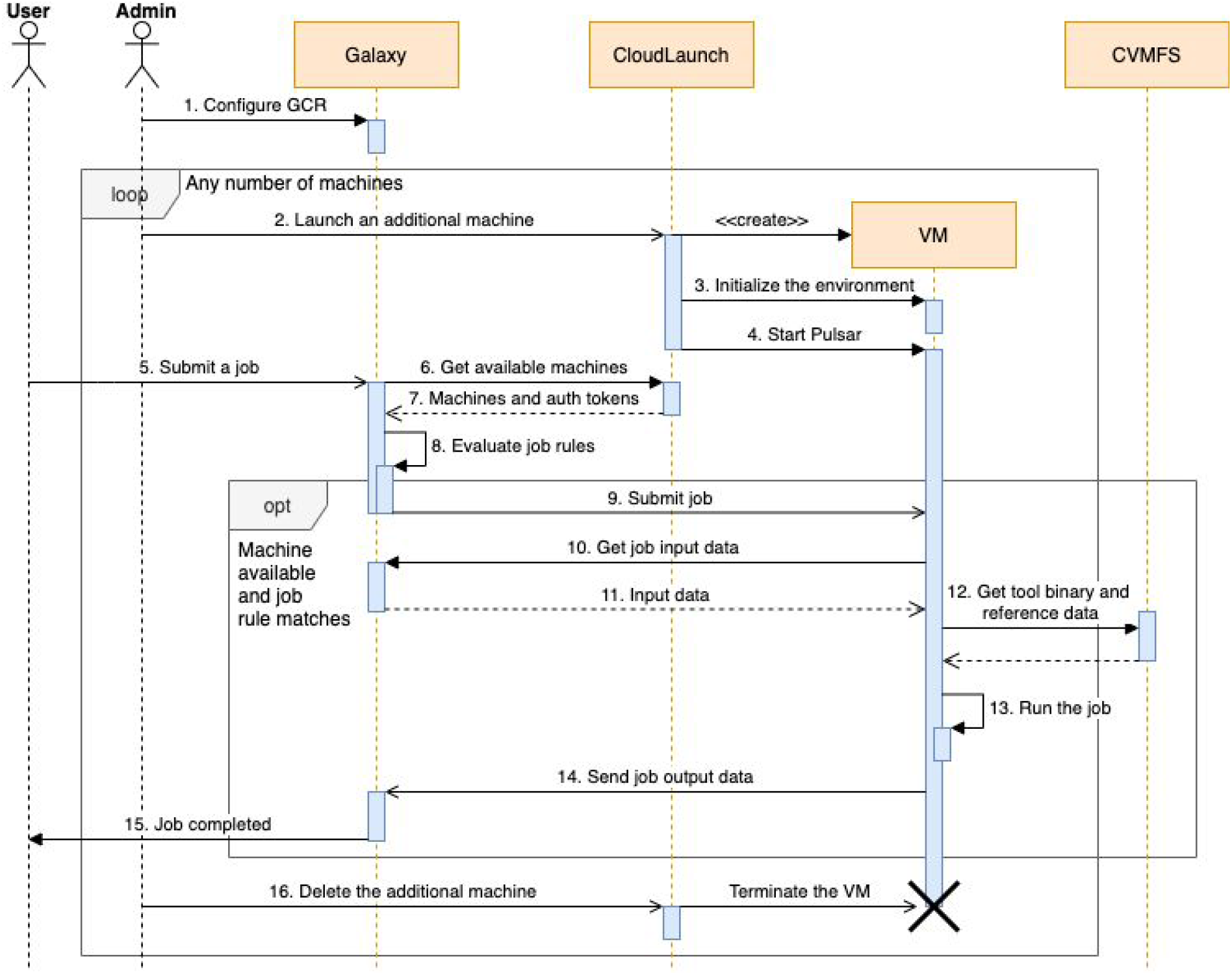
The sequence of events that occurs when bursting Galaxy jobs to the cloud via GalaxyCloudRunner (GCR). An admin enables GCR and creates any number of virtual machines at will that Galaxy will utilize based on the defined job rules. End-users use Galaxy without changing their routine.

## 3. Discussion and conclusions

GalaxyCloudRunner offers a straightforward method for acquiring additional compute capacity for any Galaxy server, does so with minimal overhead, and grants control over resources to the administrator. Suitable use cases include peak usage periods (e.g., training workshops), specific resource requirements (e.g., large memory), software licensing considerations (e.g., software available exclusively as cloud appliances), and the availability of resources from national cloud infrastructure (e.g., NSF Jetstream (Hancock *et al.*, 2019)). However, burst mode does not come without limitations. Continuously transferring large datasets from commercial cloud providers may incur sizable costs and/or take considerable time. Furthermore, not all Galaxy tools are compatible with remote execution (e.g., uploading data, data managers for reference genomes), requiring administrative care to restrict job routing in those cases. Nevertheless, the GalaxyCloudRunner is a powerful tool that can be readily used to support novel usage scenarios for Galaxy servers. For example, it is easy to envision custom job rules developed to route jobs only for specific users’ to VMs to accommodate responsive workshops or bring-your-own-resources scenarios, without needing to deploy a new Galaxy instance.

